# Response of Surface Water Quality Characteristics to Socio-economic Factors in Eastern-Central China

**DOI:** 10.1101/2021.12.19.473363

**Authors:** Maoqing Duan, Shilu Zhang, Mingxia Xu, Junyu He, Yuanyuan Gao, Jun Zhang

## Abstract

Following the implementation of the strictest water resource management system in China, it has become increasingly important to understand and improve the surface water quality and the rate at which water function zones reach the water quality standard. Based on the monthly monitoring data from 450 monitoring sites at the provincial borders of 27 provinces in China in 2019, the overall surface water quality at provincial boundaries in China was as follows: 61.7% of the water was classified under Class I–III; and 5%, 8.6%, and 12.2% of the water was classified under Class IV, V, and inferior V, respectively. The main standard items are DO, COD_Mn_, COD, BOD_5_, NH_3_-N, and TP. The Canadian Council of Ministers of the Environment-water quality index (CCME-WQI) showed that the provincial boundary water quality exceeded the fair level, and F1 was the most influential factor. Then, 27 factors that directly or indirectly affect the water quality of surface water at the provincial boundaries of 27 provinces were identified, and the indirect influencing factors were integrated into the ecological environmental quality index and human activities quantitative index. Finally, the 27 factors were integrated into six factors, and the relationship between these indicators and CCME-WQI as well as the concentration of influencing elements with respect to regulatory standard limits were analyzed. The proportion of building land was the most significant factor affecting the quality of the aquatic environment in provincial boundaries. In addition, the economic development level, proportion of farmland, and degree of social development were identified as significant influencing factors. The six factors have different degrees of impact on the concentrations of major elements with respect to standard limits. This study basically explores water resource management and offers significant reference and guidelines for the improvement of the quality of surface water at provincial boundaries in China.

## Introduction

Water resource management in China, the largest developing country in the world, regards water function zones as control units and is implementing the “most stringent water resource management system” in history[1][2]. To keep up with the developed countries with respect to water resource management, the state will examine the effectiveness of the aforementioned system based on three aspects of water quality, total water consumption, and water use efficiency for provinces. With respect to water quality, water function zones are required to record a >95% increase in water quality standard by 2030[3]. However, there is great scope for improvement of the current surface water quality status, and a lot of effort is required in this regard. The environmental water pollution events that have occurred are closely related to human activities[4]. However, the low rate of water quality standard attainment in the water function zones of certain areas (a few cases in China) is attributed to the background value rather than human activities[5]. Therefore, to improve the surface water quality in China, the rate at which water function zones reach the water quality standard should be increased, and the impact of human activities on the surface water environment reduced to meet the requirements of the “system” [6].

The continuous development and expansion of the human society and human activities have affected the surface water in different regions[7][8]. Improving the surface water environment quality and increasing the rate at which water function zones that have been adversely impacted reach the water quality standard is a top priority, determining the potential of human activities to pollute the aquatic environment, and elimination of ineffective pollution control measures are top priorities. In other words, the key surface water problems should be identified before the implementation of water environment pollution control measures, to avoid treating the symptoms rather than the root causes as the former is resource consuming and ineffective[9][10]. Therefore, the analysis of the key factors affecting the water environment quality of regional water function zones is an important reference for regional water environment treatment, and contributes to the elucidation of aquatic environment problems and avoidance of ineffective initiatives.

Based on the monthly water quality monitoring data from 450 water quality monitoring stations at the provincial boundaries of 27 provinces in China in 2019, this study 1) comprehensively analyzes and elucidates the quality of surface water at provincial boundaries in China; 2) defines the main exceed standard items that affect the quality of surface water in provincial boundaries; and 3) identifies the key factors affecting the quality of provincial boundary water bodies based on related indicators. The purpose of the study is to provide scientific reference and a theoretical basis for the targeted proposal and implementation of a strategy to improve the quality of surface water at provincial boundaries.

## Materials and Methods

### Study Area

China covers approximately 9.6 million km^2^, the surface water resources cover approximately 2.7 trillion m^3^, and the per capita occupancy is approximately 2240 m^3^[11]. The extensive utilization of water has resulted in a severe water environment crisis in China[12]. Therefore, China has initiated a stringent water resource management system, thus indicating their prioritization of the aquatic environment problem[13].

In China, the total amount and quality of surface water are managed using water function zones as control units[3]. According to the statistical data of China’s water resources annual report[14], there were 4,682 important water function zones in 2019. The study area encompassed 431 provincial boundary water function zones, 27 provinces, and 540 provincial boundary monitoring stations, as shown in Figure 1. In contrast to other types of water function zones, the quality of provincial boundary water function zones reflects the challenges of water quality in the province, and impacts the water carrying capacity of downstream provinces. Therefore, the key to avoiding conflict between provinces is to address the surface water challenges at provincial boundaries. The main control objects should be attributed to the upstream provinces.

**Fig 1.**
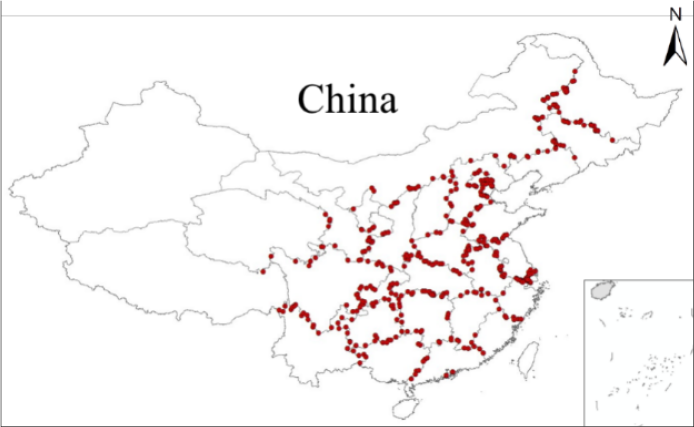
Distribution of water quality monitoring stations at provincial boundaries in 2019.

### Data Source

The upstream of the provincial boundary water quality monitoring station is regarded as the subordinate province. Table 1 presents the provincial boundary monitoring stations for each province used in this study. Data on 540 provincial boundary monitoring stations are derived from the aquatic environment monitoring centers of each watershed institution, and are monitored once a month. However, monitoring data on certain scenarios, such as dry rivers and bottom freezing, are lacking. The main monitoring items are dissolved oxygen (DO), chemical oxygen demand (COD), permanganate index (CODMn), five day biochemical oxygen demand (BOD_5_), ammonia nitrogen (NH_3_-N), total phosphorus (TP), copper (Gu), zinc (Zn), fluoride, selenium (Se), arsenic (As), mercury (Hg), cadmium (Cd), hexavalent chromium (Cr6+), lead (Pb), cyanide, volatile phenol, petroleum, anionic surfactant, and sulfide.

**Table 1.** Number of provincial boundary monitoring stations

| Province | Number | Province | Number | Province | Number | Province | Number |
| --- | --- | --- | --- | --- | --- | --- | --- |
| Beijing | 11 | Jiangsu | 52 | Hubei | 11 | Yunnan | 22 |
| Hebei | 33 | Zhejiang | 33 | Hunan | 11 | Xizang | 3 |
| Shanxi | 11 | Anhui | 20 | Guandong | 7 | Shannxi | 16 |
| Neimenggu | 38 | Fujian | 3 | Guangxi | 11 | Gansu | 14 |
| Liaoning | 5 | Jiangxi | 5 | Chongqing | 8 | Qinghai | 9 |
| Jilin | 14 | Shandong | 20 | Sichuan | 19 | Ningxia | 6 |
| Heilongjiang | 19 | Henan | 26 | Guizhou | 23 | / | / |

Considering the natural and anthropogenic factors affecting surface water, 27 factors that directly or indirectly affect the surface water quality were identified from the 2019 China Environment Statistical Yearbook[15] and the Resource and Environment Science and Data Center of the Chinese Academy of Sciences[16], as shown in Table 2.

**Table 2.** Factors influencing the quality of provincial boundary water bodies

| Number | Indicators | Unit | Number | Indicators | Unit |
| --- | --- | --- | --- | --- | --- |
| 1 | Farmland | km<sup>2</sup> | 15 | Economic density | yuan×10<sup>4</sup>/km<sup>2</sup> |
| 2 | Woodland |  | 16 | Agricultural output value per unit area |  |
| 3 | Grassland |  | 17 | Forestry output value per unit area |  |
| 4 | Building land |  | 18 | Output value of animal husbandry per unit area |  |
| 5 | Unused land |  | 19 | Fixed assets investment per unit area |  |
| 6 | Water body |  | 20 | Total power of agricultural machinery per unit area | kW/km<sup>2</sup> |
| 7 | Surface water resources per unit area (Qs) | m<sup>3</sup>/km<sup>2</sup> | 21 | Per capita output of main agricultural products | kg |
| 8 | Population density | person/km<sup>2</sup> | 22 | Total livestock per unit area | /km<sup>2</sup> |
| 9 | Proportion of urban population | % | 23 | Number of secondary school students per 100000 population | person |
| 10 | Proportion of employees in the whole society |  | 24 | Number of ordinary secondary schools per unit area | per/km<sup>2</sup> |
| 11 | Road density | / | 25 | Number of teachers per unit area | person/km<sup>2</sup> |
| 12 | Natural population growth rate |  | 26 | Total wastewater discharge per unit area (Qw) | t/km<sup>2</sup> |
| 13 | Per capita gross national product | Yuan | 27 | Unit domestic waste removal and transportation |  |
| 14 | Application rate of chemical fertilizer per unit | t/km<sup>2</sup> | / | / | / |

Factors that can directly and evidently pollute the surface water environment are the application rate of chemical fertilizer per unit, proportion of farmland, proportion of wastewater discharge in the total surface water resources, and unit domestic waste removal and transportation. The remaining factors were organized into a comprehensive index based on ecological environmental quality and human activity intensity.

## Evaluation Method

### Single factor evaluation method

According to Environmental Quality Standards for Surface Water (GB2002-3838)[17], the single factor evaluation method was used to classify the water quality monitoring items into different water quality categories (I-V Class) with different concentration values.

### Comprehensive index score method

CCME-WQI is a method that accounts for all variables and yields a single value based on the classification by the CCME[18]. The index comprises three factors: Factor 1 (F_1_) is the percentage of the variables exceeding allowable limits; Factor 2 (F_2_) is the percentage of the quality of samples exceeding allowable limits during the study; and Factor 3 (F_3_) is the amplitude exceeding the standard[19] [20]. These factors are expressed as follows:.

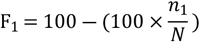

where N is the total number of variables and n_1_ is the number of variables whose values do not exceed the standard during the monitoring period.

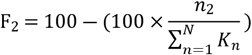

where N is the number of monitoring variables, n_2_ is the number of times the monitoring variable does not exceed the standard during the monitoring period, and K_n_ is the total monitoring frequency of the variable.

F_3_ was calculated in two steps. When the monitoring value of a variable failed to meet the objective and was below the standard value, the following equation applied:

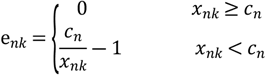

Conversely, when the monitoring value of a variable failed to meet the objective and exceeded the standard value, the following equation was applied:

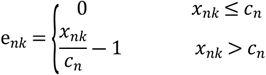

where enk is the deviation of the variable, xnk is the monitoring value of the variable, and cn is the standard value.

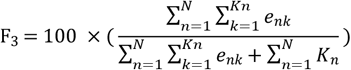

The values of F_3_ ranged from 0 to 100 and reflected the amplitude by which the variable exceeded the standard. Finally, the CCME-WQI was calculated as follows:

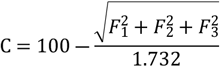

The following five levels of water quality were then determined:

1. Excellent: (CCME WQI ranging from 95–100);
2. Good: (CCME WQI ranging from 80–94);
3. Fair: (CCME WQI ranging from 60–79);
4. Marginal: (CCME WQI ranging from 45–59);
5. Poor: (CCME WQI ranging from 0–44).

### Ecological Environmental Quality Index

EEQI is calculated using the proportion of land use area and represents the ecological quality index as follows:

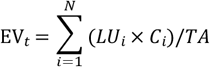

EVt- EEQI; LUi-Area of land use type i: Ci-Ecological quality index of land use type i (Table 3); N-Land use types; TA-Total area.

**Table 3.** Ecological quality index of various land use types

| Primary type |  | Secondary type |  | Ecological quality index | Primary type |  | Secondary type |  | Ecological quality index |
| --- | --- | --- | --- | --- | --- | --- | --- | --- | --- |
| Number | Type | Number | Type |  | Number | Type | Number | Type |  |
| 1 | Farmland | 11 | Paddy field | 0.3 | 4 | Water body | 41 | Rivers and canals | 0.55 |
|  |  | 12 | Dry land | 0.25 |  |  | 42 | Lakes | 0.75 |
| 2 | Woodland | 21 | Forest land | 0.95 |  |  | 43 | Reservoirs | 0.55 |
|  |  | 22 | Shrub wood | 0.65 |  |  | 44 | Permanent River Snow | 0.9 |
|  |  | 23 | Sparse Woodland | 0.45 |  |  | 45 | Beach | 0.45 |
|  |  | 24 | Others | 0.4 |  |  | 46 | Wetlands | 0.55 |
| 3 | Grassland | 31 | High coverage | 0.75 | 6 | Unused land | 61 | Sand | 0.01 |
|  |  | 32 | Medium coverage | 0.45 |  |  | 62 | Gobi | 0.01 |
|  |  | 33 | Low coverage | 0.2 |  |  | 63 | Saline-alkali soil | 0.05 |
| 5 | Building land | 51 | Urban land | 0.2 |  |  | 64 | Swamp | 0.65 |
|  |  | 52 | Rural land | 0.2 |  |  | 65 | Bare land | 0.05 |
|  |  | 53 | Others | 0.15 |  |  | 66 | Bare rock gravel | 0.01 |

### Ecological Environmental Quality Index

The human activities quantitative model and index weight value were proposed by Zhigang Xu[21]. The specific steps of measuring regional human activity intensity from three aspects: society, economy, and culture are as follows: 1) identification of the dimensionless index, 2) calculation of index weight value, and 3) determination of the regional human activity intensity index by index weighting. The evaluation index system is shown in Figure 3.

## Result

### Water Quality Analysis

According to the Surface Water Environmental Quality Standard (GB3838-2002)[17], the standard of provincial boundary water quality falls under class II–V. The eigenvalues of the 20 monitoring items of provincial boundary water quality in China in 2019 are shown in Table 4. Overall, the average concentrations of TP fall under Class III, COD_Mn_ and NH_3_-N under Class II, and the other elements under Class I. Gu, Zn, Se, As, cyanide, petroleum, and sulfide at concentrations are relatively low. The quality standards of the maximum concentration values monitored in 2019 were below Class V. The maximum concentrations of volatile phenol, Cr^6+^, COD_Mn_, fluoride, Cd, BOD_5_, anionic surfactant, and COD were 1.37–3.7 times higher than the Class V standard limits, and TP and NH_3_-N exceeded these limits by 20.1 and 14.9 times, respectively. In 2019, 540 provincial boundaries were monitored 4,910 times, with Class II water quality accounting for the highest proportion (32.3%), followed by Class III water quality (22.9%). Class I water quality accounted for the lowest proportion (6.5%). The proportion of inferior Class V water quality was relatively high, accounting for 12.2%. Therefore, additional attention should be paid to water quality control for stations with poor water quality (Class **V**).

**Table 4.**
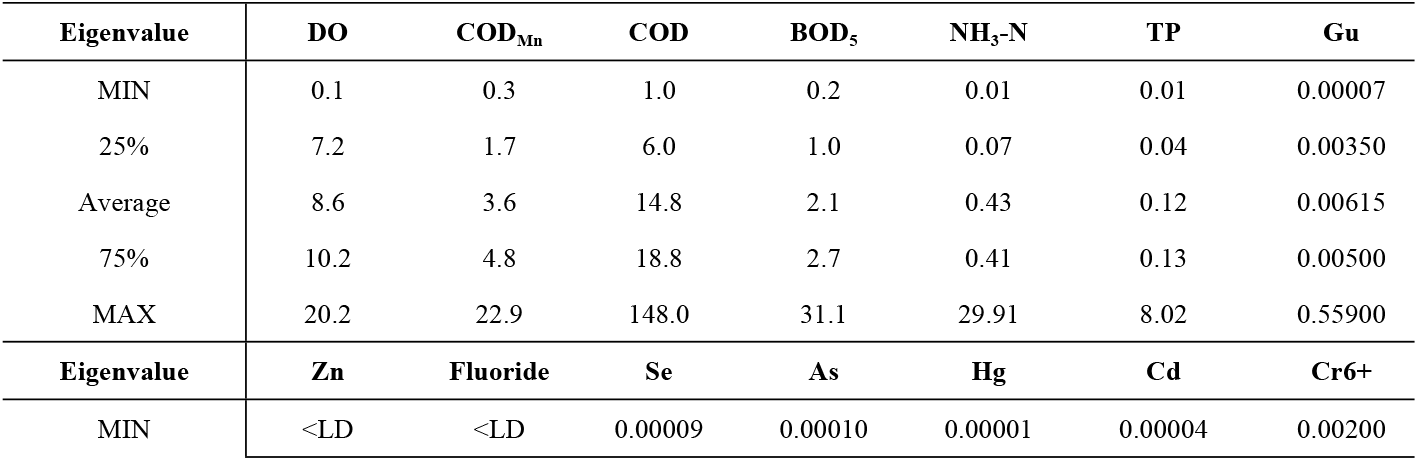
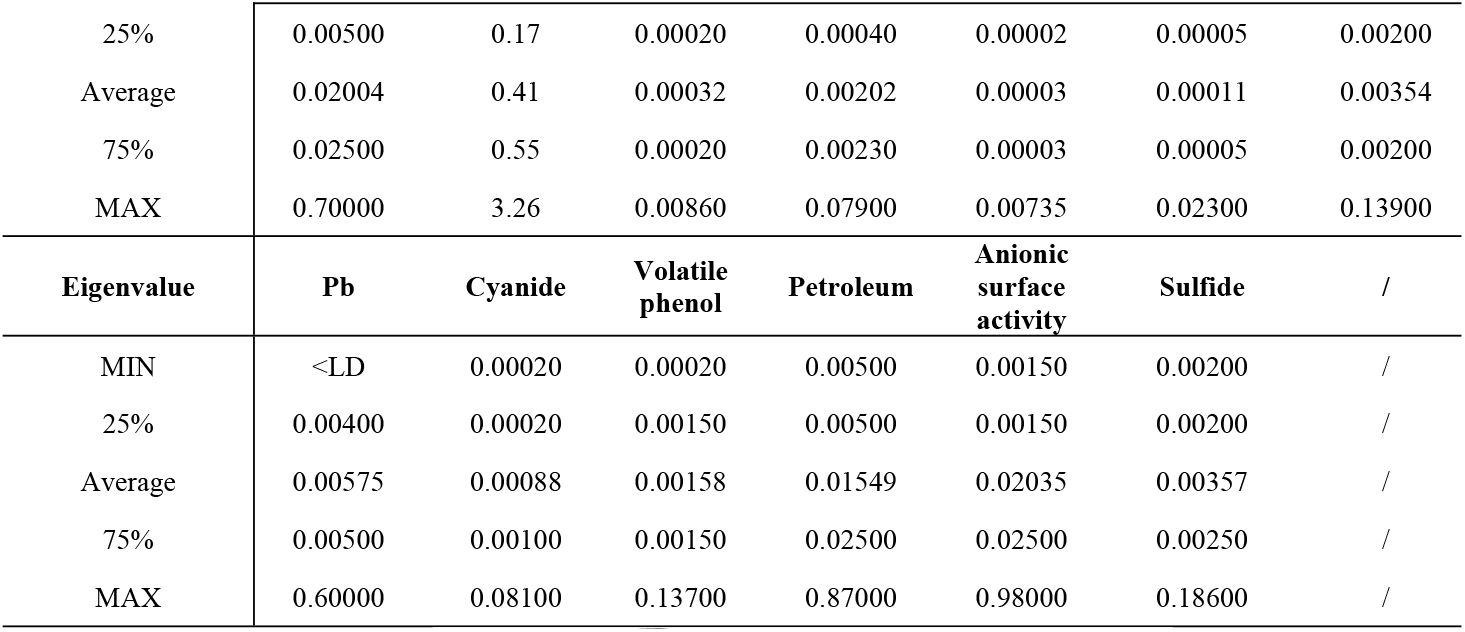
Eigenvalue concentration value of water quality items in 2019 (mg/L)

The 540 provincial boundary stations implement different water quality standards. As seen in Figure 2, COD and TP are the main elements exceeding standard limits, with the exceeding rates of 18.4% and 12.3%, respectively, followed by NH_3_-N, COD_Mn_, and BOD_5_, with exceeding rates of 9%, 8.3%, and 8.6%, respectively. DO and fluoride exceed standard limits by 5.8% and 3.4%, respectively, and other elements by less than 2%. The exceeding standard rates of Fluoride, Petroleum, Cr^6+^, Hg, Zn, Se, Cyanide, Sulfide, Volatile phenol, Pb, Anionic surface activity, Cd, As, and Gu are below 5%. There are only less than 170 times exceeding standard in the more than 4000 times detection, indicating that the remaining 14 items have no significant impact on the quality of surface water in China’s provincial boundaries, and therefore do not require much attention. However, COD,TP,NH_3_-N,COD_Mn_,BOD_5_, and DO exceed standard limits by more than 5%, and significantly impact the quality of surface water at provincial boundaries in China, and hence require attention.

**Fig 2.**
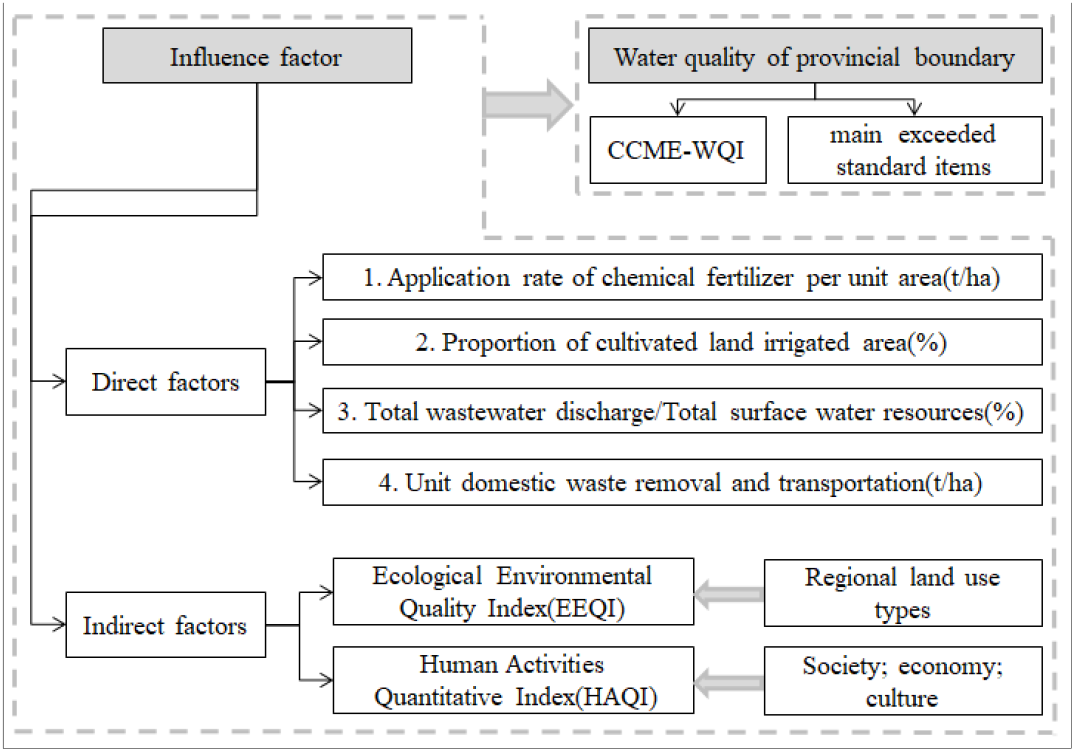
Framework for analysis of factors influencing the quality of provincial boundary waterbodies

**Fig 3.**
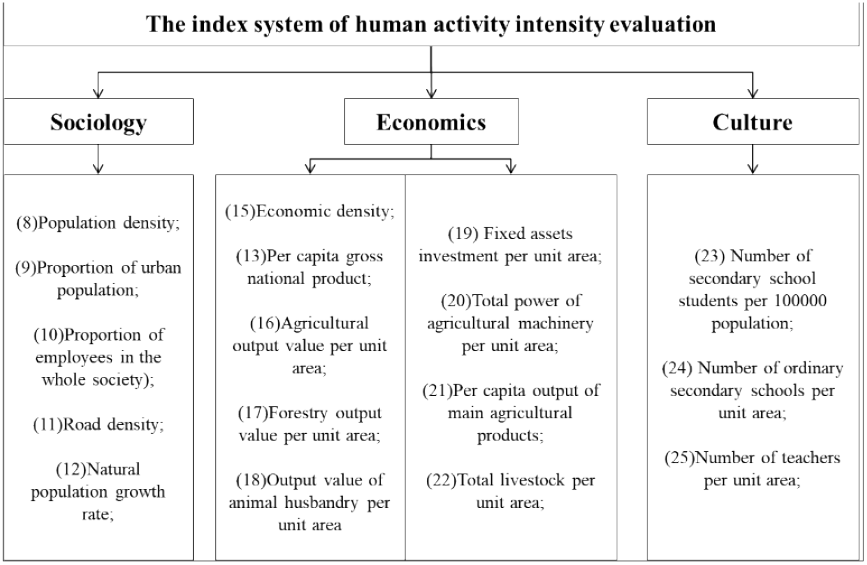
Quantitative index system of regional human activity intensity

**Fig 4.**
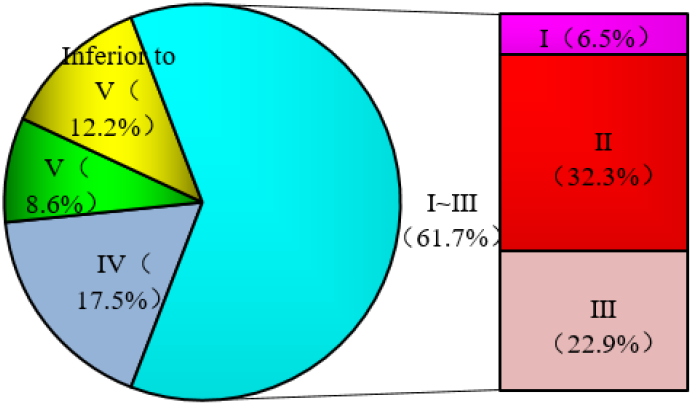
Proportion of water quality categories at provincial boundaries in 2019

**Fig 5.**
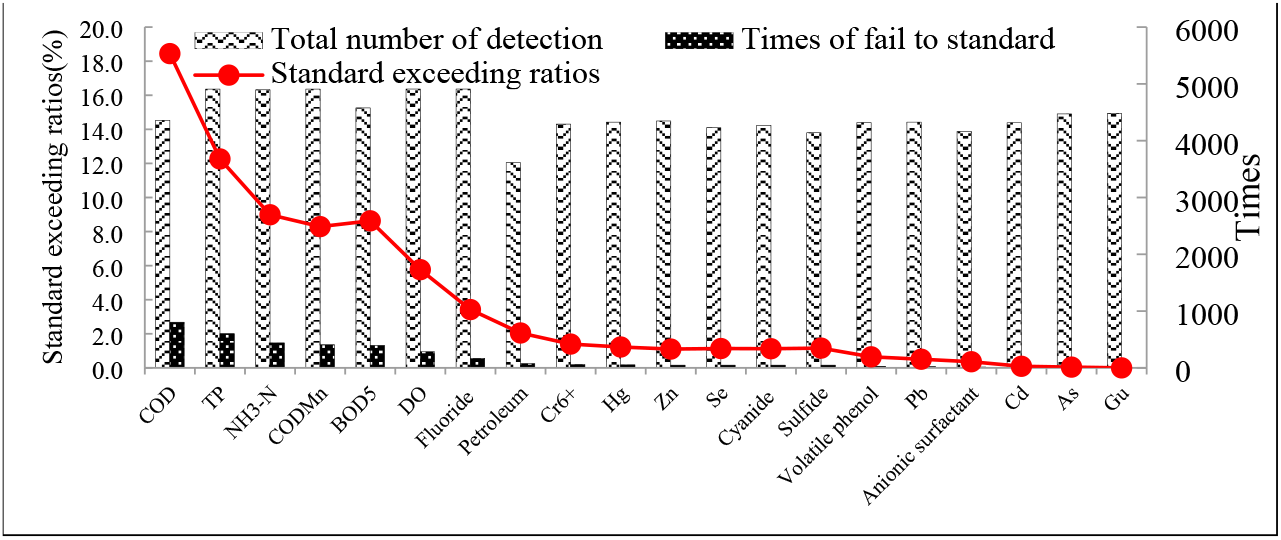
Monitoring frequency, and exceeding rates of each item

The provincial boundary water quality monitoring stations are regarded as the representative stations of the upstream cities or provinces and reflect the distribution of the main elements standard limits (Figure 6). The exceeding standard of COD, which reflects the total amount of oxygen-consuming organic matter in the water body, is the highest: 50.3% in Hebei, 46.4% in Henan, and more that 20% in Neimeng, Jilin, Heilongjiang, Anhui, and Shandong. In addition, the exceeding COD rates are dominant in Central, East and Northeast China. Phosphorus is an important nutrient for water life; however, excess phosphorous causes rapid reproduction of algae and other plankton, decrease in dissolved oxygen content, and water quality deterioration[22]. TP in Liaoning, Shandong, Jilin, Henan, and Neimeng exceeded the standard limit by 20%, which was lower than the exceeding rate of COD and exhibited a similar spatial distribution in contrast with that of COD. NH_3_-N can be converted into nitrite under certain conditions in water, and has adverse effects on human health and aquatic life[23]. The highest rate of NH_3_-N exceeding the standard limit in Guangdong Province was 29.6%, the second highest was Beijing (28.6%), and provinces with high exceeding rates with respect to standard limits are mainly concentrated in the central and eastern regions. The spatial distribution of COD_Mn_ exceeding the standard limit is similar to that of COD and TP; Heilongjiang, Jilin, and Hebei reports a 20% exceeding rate, and higher exceeding rates are dominant in the Middle East and northeast regions. BOD_5_ reflects the total amount of organic matter that can be decomposed by microorganisms in water[24], Zhejiang reports the highest exceeding standard rate (21.5%), followed by Jiangsu (19.8%) and Guangdong (17.3%). Finally, Zhejiang, Jiangsu, and Guangdong are still found in provinces with an exceeding rate of more than 10%, and the highest was in Fujian Province.

**Fig 6.**
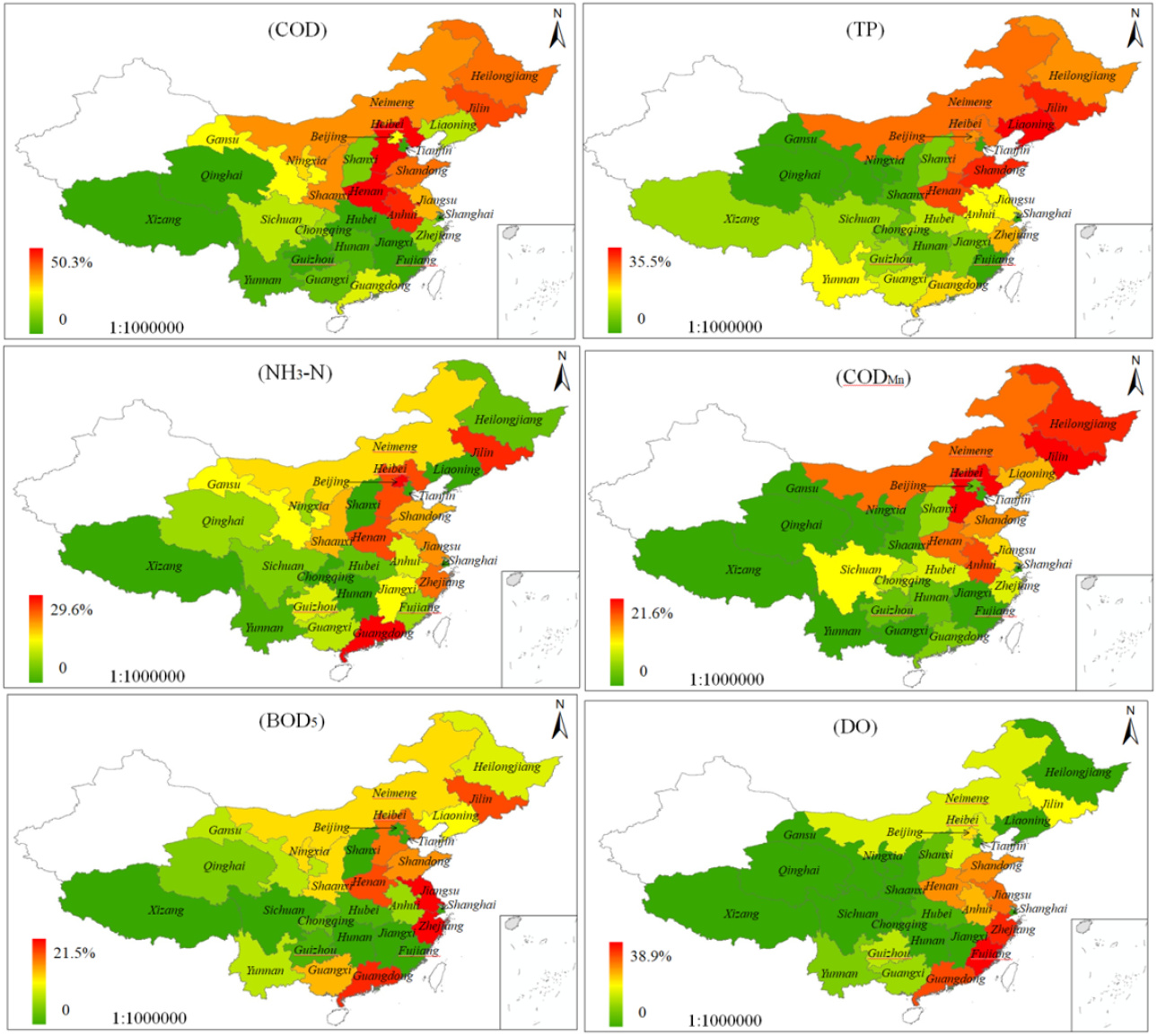
Spatial distribution of main elements exceeding standard limits

From the spatial distribution of the main elements exceeding the standard limits, oxygen consumption, and nutrients were mainly concentrated in the Middle East Northeast China. BOD_5_ and DO mainly exceeded the standard limits in the southeast coastal area. In the central and southwest regions of China, the exceeding rate of water quality elements was relatively low. This phenomenon is related to the level of economic development and the spatial distribution of human social development.

### Canadian Council of Ministers of the Environment water quality index

The composite index score (Figure 7) of CCME-WQI shows that Xizang, which yielded the highest score (97.2), is at the excellent level, Jiangxi to Anhui are at the good level, Yunnan to Hebei are at the fair level, and Neimeng are at the marginal level. There were no provincial boundary water quality scores at the poor level, the national provincial water quality score is 82.1, which is at a remarkable level, and the water quality is generally good. F1 contributes the most to C. Many elements exceed standard limits according to water quality monitoring data. However, F2 shows that the total exceeding rate is very low, Shandong, Guangdong, Jilin, Heibei, Henan report higher rates of exceeding standard amplitude limits (F3), and exceeding rates of other elements are below 10%.

**Fig 7.**
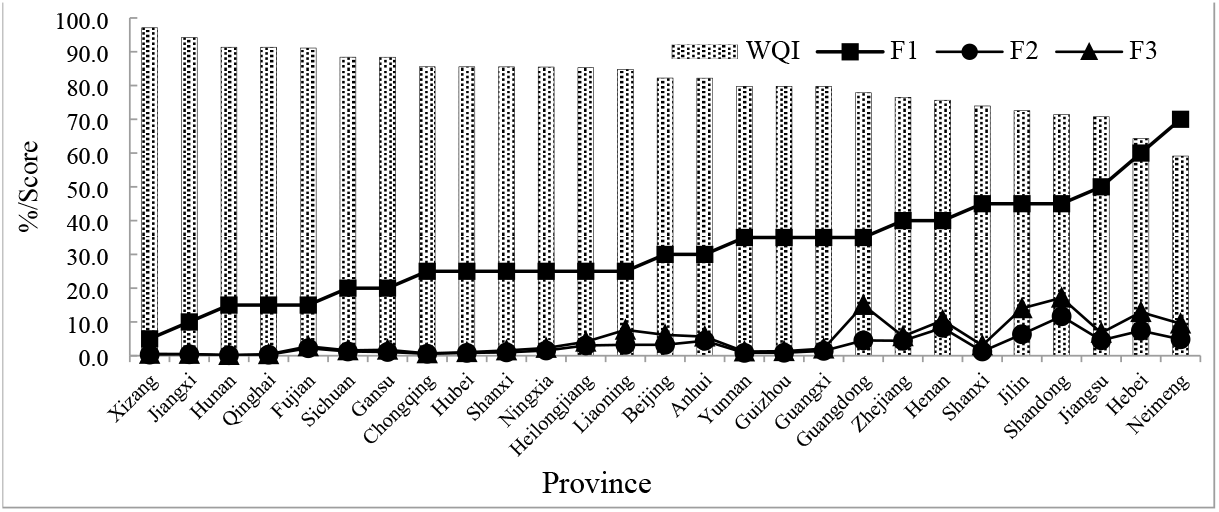
Comprehensive index scores of water quality at provincial boundaries

Figure 8 is plotted according to F2. There are six main elements exceeding standard limits, COD and TP exceeded the standard limit in 18 provinces, followed by COD_Mn_ and NH_3_-N, which exceeded the standard limit in 12 provinces. In contrast, DO exceeded the standard limits in only four provinces. Therefore, COD and TP exceeded standard limits most frequently at provincial boundaries in China, and are the essential elements during the aquatic environment control at provincial boundaries.

**Fig 8.**
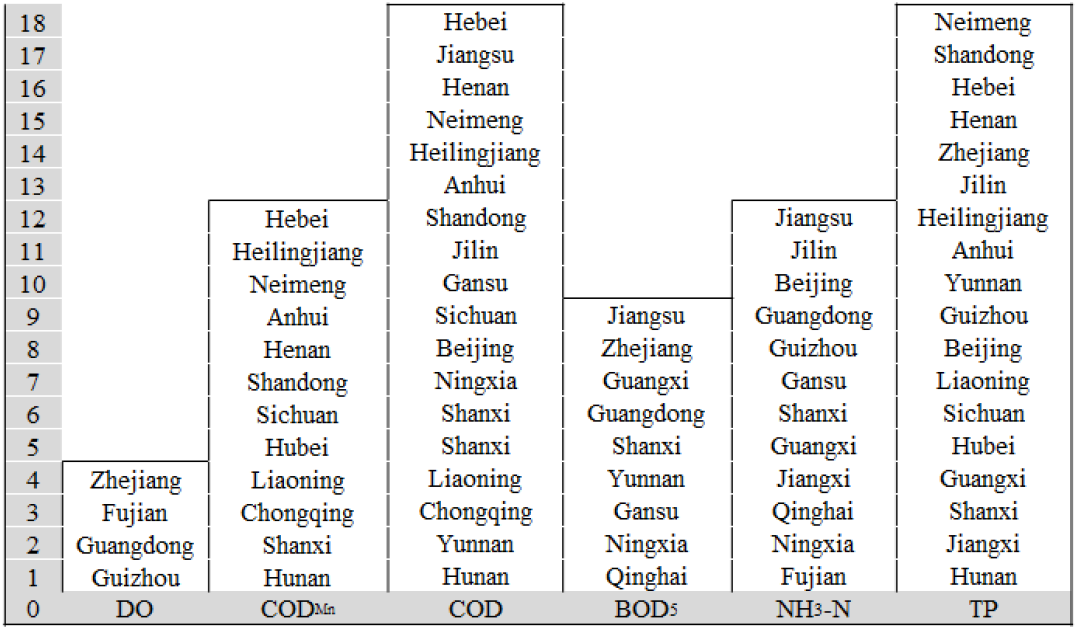
Provinces where six elements exceeded standard limits (The ordinate represents the ranking of F2 values in different provinces)

Hebei province should focus on COD_Mn_, COD, and TP (COD_Mn_ and COD exceeded standard limits most frequently here). Second, Jiangsu province should focus on BOD_5_, NH_3_-N, and COD (BOD_5_ and NH_3_-N as these exceeded standard limits most frequently followed by COD. Finally, Neimeng should focus on TP, COD, and COD_Mn_.

### Factors directly influencing surface water quality

Among the 27 factors identified, the application rate of chemical fertilizer per unit area (t/ha), proportion of cultivated and irrigated area (%), total wastewater discharge/total surface water resources (%), and unit domestic waste removal and transportation (t/ha) were identified as having the capacity to directly pollute the surface water environment and were analyzed (Figure 9). The application rate of chemical fertilizer per unit area ranged from 0.04 to 41.5 t/km^2^, and those of Henan, Jiangsu, Shandong, Anhui exceeded 20 t/km^2^. Unit domestic waste removal and transportation ranged from 0.4 to 580.4 t/km^2^, the maximum values were yielded in Beijing, Guangdong, Jiangsu, Zhejiang, Shandong which exceeded 100 t/km^2^. Qw/Qs values ranged from 0.02–93.1, and those yielded by Beijing, Hebei, Ningxia, Shandong, Jiangsu exceeded 20%. The proportion of cultivated and irrigated area ranged from 0.2–40.9%, and Jiangsu, Anhui, Shandong, Hebei, Henan reported over 20%, Overall, the four factors mentioned above show that the eastern region is larger than the western region, and the northern region is larger than the southern region.

**Fig 9.**
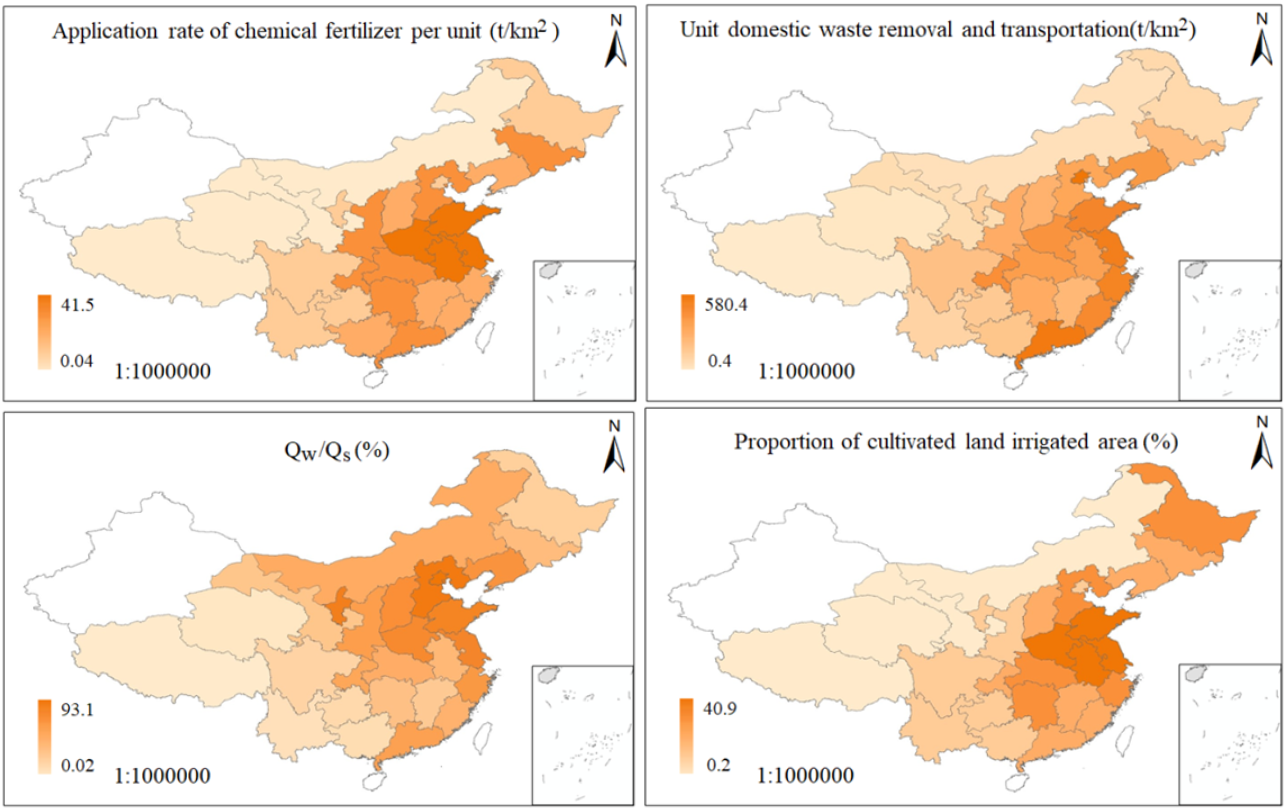
Spatial distribution map of factors directly affecting water quality

**Fig 10.**
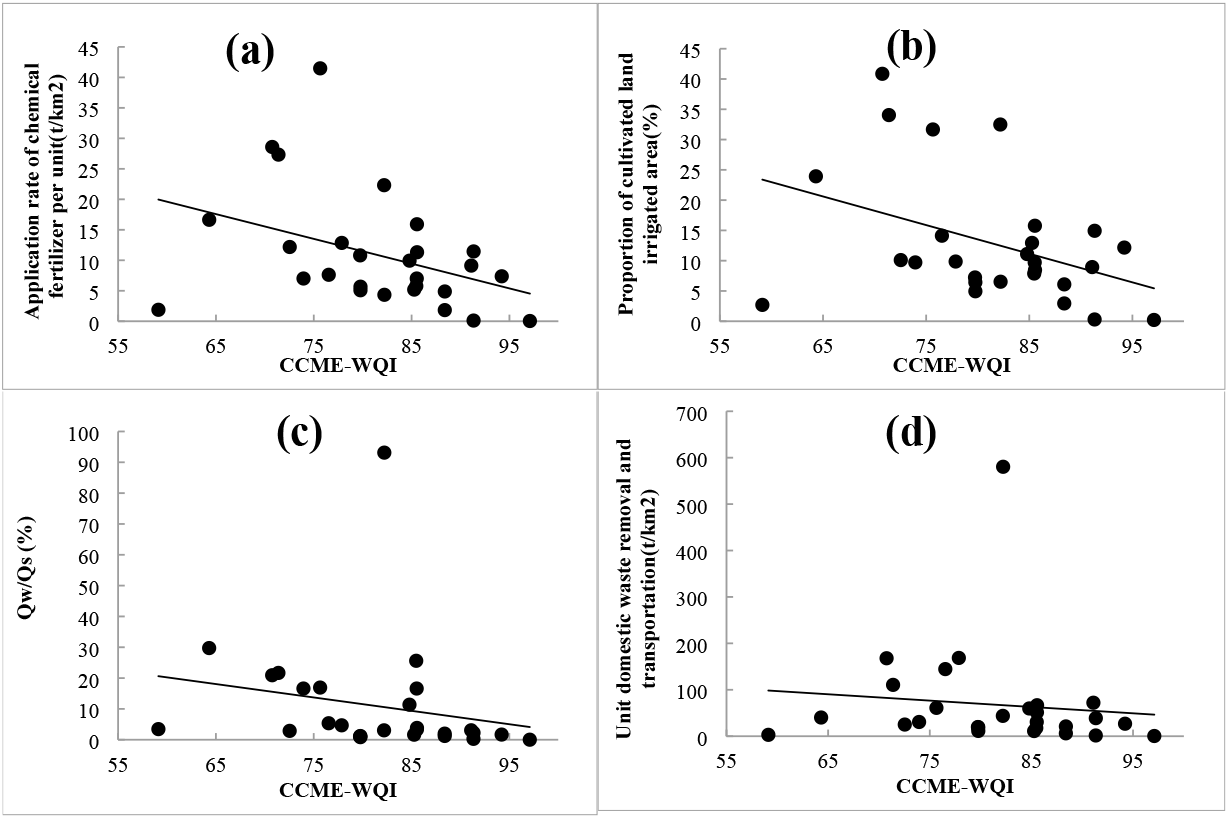
Correlation analysis between water quality and direct influencing factors ((a)-application rate of chemical fertilizer per unit area (t/ha);(b)-Proportion of cultivated land irrigated area (%); (c)-total wastewater discharge/Total surface water resources (%); (d)-unit domestic waste removal and transportation (t/ha))

### Factors indirectly influencing surface water quality

To reflect their impact on provincial boundary water quality, factors that indirectly influence surface water quality can be divided into two: natural (EEQI) and anthropogenic (HAQI) factors.

#### 1) Environmental Ecological Quality Index

Land use is the main factor influencing non-point sources, and different land use patterns lead to spatial differences in the degree and spatial distribution of pollution[25]. Among the 27 provinces, the proportion of farmland ranged from 0.6% to 63.9%, with Shandong accounting for the largest proportion, followed by Henan (62.5%). Agricultural production is an important tool of controlling non-point source pollution. The types of cultivated land, farming methods, and crop types have obvious effects on the non-point source pollution. Woodland accounts for 3–65.5% of land use, the largest proportion is Guangxi. Grassland accounts for 1.1-55.5% of land use, the largest being in Qinghai and the smallest in Jiangsu. Building land indirectly reflects the intensity of regional human activities. Among the 27 provinces, building land accounts for 0.1–21.3%, the largest being in Jiangsu, followed by Beijing. Waterbodies account for 0.6–14% of land use, the largest being Jiangsu, the proportion of unused land is 0–37.8%, and the largest proportion is in Gansu. The EEQI of 27 provinces were calculated based on land use type and their ecological quality index values shown in Table 3. The results are presented in Table 5.

**Table 5.**
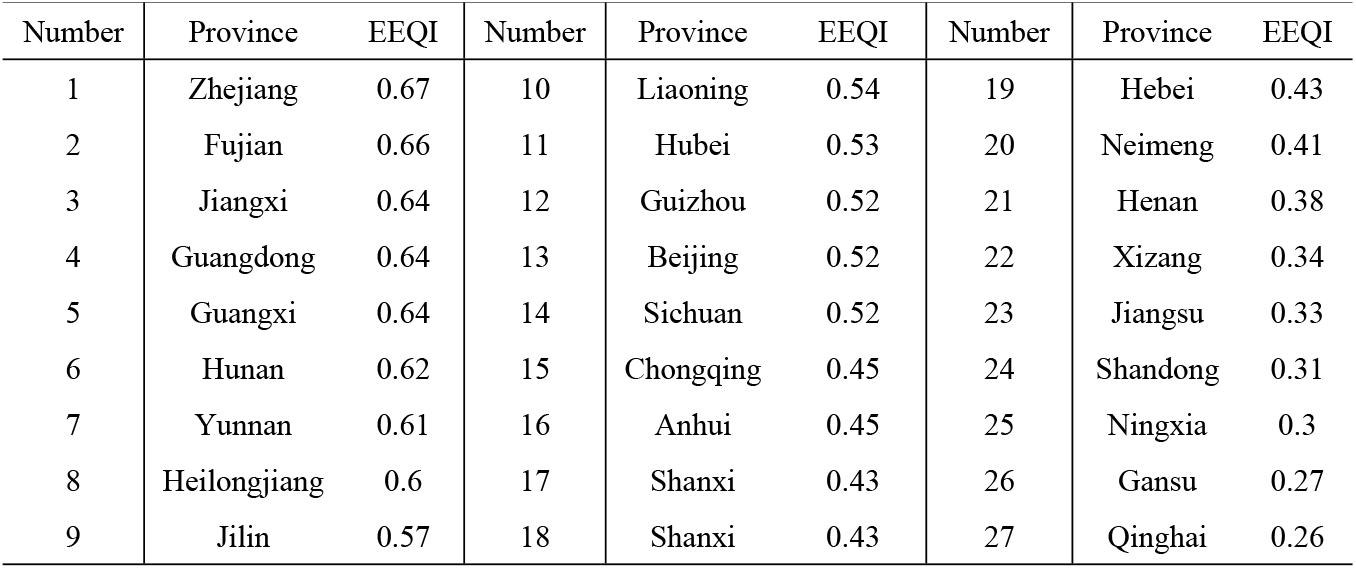
Ecological environmental quality index of 27 provinces

| Number | Province | EEQI | Number | Province | EEQI | Number | Province | EEQI |
| --- | --- | --- | --- | --- | --- | --- | --- | --- |
| 1 | Zhejiang | 0.67 | 10 | Liaoning | 0.54 | 19 | Hebei | 0.43 |
| 2 | Fujian | 0.66 | 11 | Hubei | 0.53 | 20 | Neimeng | 0.41 |
| 3 | Jiangxi | 0.64 | 12 | Guizhou | 0.52 | 21 | Henan | 0.38 |
| 4 | Guangdong | 0.64 | 13 | Beijing | 0.52 | 22 | Xizang | 0.34 |
| 5 | Guangxi | 0.64 | 14 | Sichuan | 0.52 | 23 | Jiangsu | 0.33 |
| 6 | Hunan | 0.62 | 15 | Chongqing | 0.45 | 24 | Shandong | 0.31 |
| 7 | Yunnan | 0.61 | 16 | Anhui | 0.45 | 25 | Ningxia | 0.3 |
| 8 | Heilongjiang | 0.6 | 17 | Shanxi | 0.43 | 26 | Gansu | 0.27 |
| 9 | Jilin | 0.57 | 18 | Shanxi | 0.43 | 27 | Qinghai | 0.26 |

#### 2) Human activities quantitative index

Human activities are the driving factors influencing the surface water quality, and the quantitative evaluation of regional human activity intensity is the basis for analyzing the impact of human activities on surface water environmental quality. Because of the uncertainty and complexity of human activities, it is difficult to quantitatively evaluate regional human activities. Here, we refer to studies that established the Regional Quantitative Model of Human Activity Intensity to calculate the human activities quantitative index (HAQI) of 27 provinces (Table 6); Jiangsu yielded the highest value (0.56); followed by Shandong (0.52); the minimum value was reported by Xizang (0.08). From the contribution of various aspects, the contribution of cultural factors to HAQI is the lowest, all below 0.04 in 27 provinces, followed by economic factors, and the highest value is 0.13 in Shandong. The social factor is the main contributor to HAQI.

**Table 6.** Quantitative index of human activities in 27 provinces of China(HAQI)

| Province | Society | Economy | Culture | Composite index | Province | Society | Economy | Culture | Composite index |
| --- | --- | --- | --- | --- | --- | --- | --- | --- | --- |
| Beijing | 0.34 | 0.08 | 0.02 | 0.44 | Hubei | 0.17 | 0.06 | 0.02 | 0.25 |
| Hebei | 0.2 | 0.07 | 0.03 | 0.30 | Hunan | 0.16 | 0.06 | 0.03 | 0.25 |
| Shanxi | 0.13 | 0.02 | 0.02 | 0.17 | Guangdong | 0.26 | 0.06 | 0.03 | 0.35 |
| Neimeng | 0.07 | 0.03 | 0.01 | 0.11 | Guangxi | 0.13 | 0.04 | 0.04 | 0.21 |
| Liaoning | 0.15 | 0.05 | 0.01 | 0.21 | Chongqing | 0.18 | 0.05 | 0.03 | 0.26 |
| Jilin | 0.09 | 0.05 | 0.01 | 0.15 | Sichuan | 0.09 | 0.03 | 0.02 | 0.14 |
| Heilongjiang | 0.07 | 0.04 | 0.01 | 0.12 | Guizhou | 0.10 | 0.04 | 0.05 | 0.19 |
| Jiangsu | 0.42 | 0.12 | 0.02 | 0.56 | Yunnan | 0.08 | 0.03 | 0.03 | 0.14 |
| Zhejiang | 0.28 | 0.06 | 0.02 | 0.36 | Xizang | 0.04 | 0.01 | 0.03 | 0.08 |
| Anhui | 0.24 | 0.08 | 0.03 | 0.35 | Shaanxi | 0.12 | 0.03 | 0.02 | 0.17 |
| Fujian | 0.21 | 0.05 | 0.03 | 0.29 | Gansu | 0.06 | 0.01 | 0.02 | 0.09 |
| Jiangxi | 0.16 | 0.04 | 0.04 | 0.24 | Qinghai | 0.07 | 0.01 | 0.03 | 0.11 |
| Shandong | 0.36 | 0.13 | 0.03 | 0.52 | Ningxia | 0.11 | 0.03 | 0.04 | 0.18 |
| Henan | 0.21 | 0.11 | 0.04 | 0.36 | / | / | / | / | / |

**Table 7.**
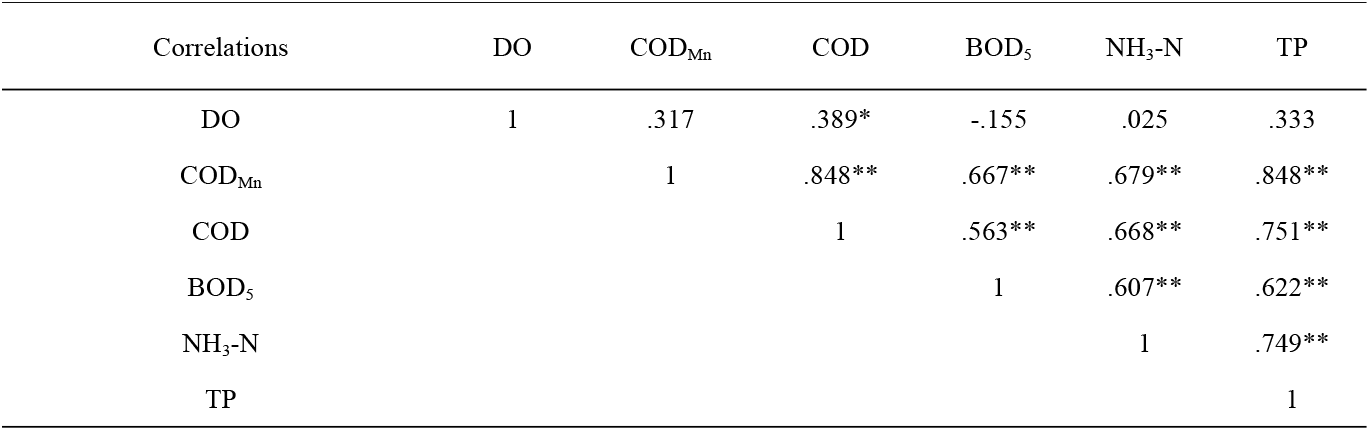
Pearson correlation analysis of major elements exceeding the standard limits

| Correlations | DO | $\text{COD}_{\text{Mn}}$ | COD | $\text{BOD}_5$ | $\text{NH}_3\text{-N}$ | TP |
| --- | --- | --- | --- | --- | --- | --- |
| DO | 1 | .317 | .389* | -.155 | .025 | .333 |
| $\text{COD}_{\text{Mn}}$ | | 1 | .848** | .667** | .679** | .848** |
| COD |  |  | 1 | .563** | .668** | .751** |
| $\text{BOD}_5$ | | | | 1 | .607** | .622** |
| $\text{NH}_3\text{-N}$ | | | | | 1 | .749** |
| TP |  |  |  |  |  | 1 |

## Discussion

### Factors Influencing Water Quality

The natural environment and human activities are the two major factors affecting water quality; however, the complexity and inherent interdisciplinary characteristics of the ecological environment determine the difficulty of comprehensive evaluation[26][27]. In this study, 27 factors affecting water environment quality were identified to represent the natural environment and human activities, and the indirect influencing factors were expressed in the form of a comprehensive index to reflect the impact of ecological environment quality and human activity intensity on water quality.

We analyzed the relationship between the CCME-WQI and the four direct influencing factors, and found a negative but insignificant correlation between the water quality, and application rate of chemical fertilizer per unit area (R^2^=0.15) and the proportion of cultivated land irrigated area (R^2^=0.16) at provincial boundaries. There was no correlation between the water quality and total wastewater discharge/total surface water resources, unit domestic waste removal, and transportation (R^2^<0.05). In the case where point source pollution is gradually controlled, agricultural non-point source pollution is the main driving factor of surface water quality change at provincial boundaries.

Figure 11(a) shows that there was no correlation between the CCME-WQI and the EEQI (R^2^=0.0112), which means that the regional eco-environmental quality is not the main control factor affecting the provincial water quality. However, we found that the chemical fertilizer per unit area and proportion of cultivated land irrigated area have a relatively good correlation with water quality. The calculation results of the EEQI may conceal the impact of various land use types on the water environment quality, so we analyzed the relationship between the six types of land use and CCME-WQI. The results yielded R^2^ values of 0.16 for farmland, 0.22 for building land, and 0.05 for the other four factors. Therefore, human activities are evidently the main influencing factors of provincial water quality, followed by agricultural non-point source pollution.

**Fig 11.**
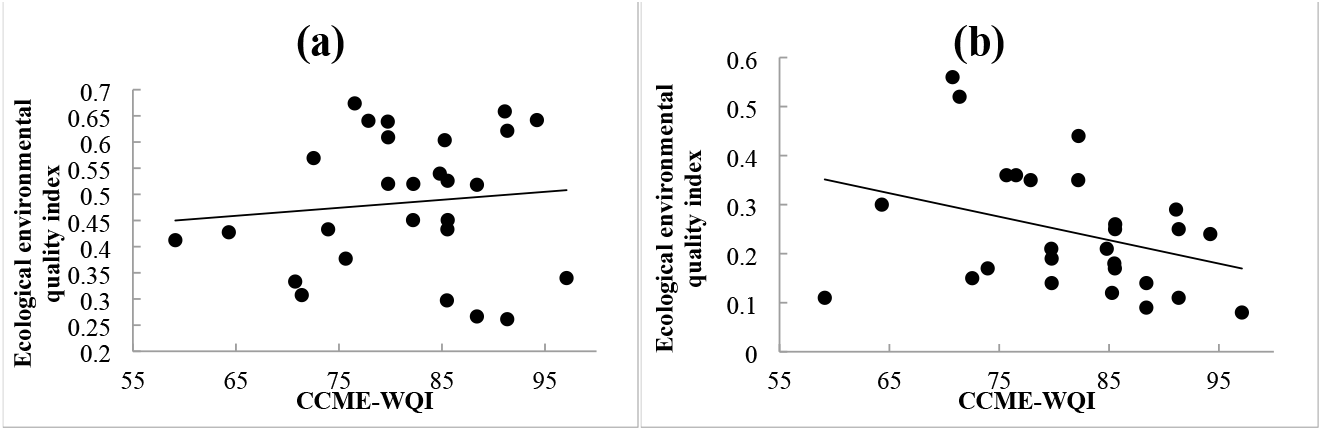
Correlation analysis between water quality and composite index (a-ecological environmental quality index; b-human activities quantitative index)

Furthermore, upon analyzing the relationship between the HAQI and the CCME-WQI, we determined R^2^ to be 0.1144 and to have a more significant impact than EEQI (Figure 11b). Considering the impact of social, economic, and cultural aspects on the aquatic environment, the correlation values between these aspects and water quality were 0.1044, 0.1877, and 0.05, respectively. Therefore, the intensity index of human activities and economic development is the controlling factor of water quality, followed by the degree of societal development.

According to the analysis results of six indicators and the CCME-WQI, human activities are the important factors affecting the water quality at provincial boundaries. The proportion of building land was the most important factor (R^2^=0.22), followed by the development of the regional social economy (0.1877), the development of farmland (0.15 ∼ 0.16), and the degree of regional social development (0.1044), respectively.

### Factors influencing element concentration

In addition to DO, there is a significant correlation between the concentrations of five elements with evident homology exceeding the standard limits, and their correlation coefficients range from 0.56–0.84 (P < 0.01). Table 8 shows the relationship between the main elements exceeding the standard limits and the six main influencing factors. COD_Mn_ and COD exhibited the lowest correlation with EEQI; relative to the other five factors. BOD_5_ exhibited a strong correlation with the application rate of chemical fertilizer per unit, proportion of cultivated land irrigated area, and HAQI. NH_3_-N exhibited a strong correlation to Qw/Qs, unit domestic waste removal and transportation, and HAQI. There was no correlation between TP and EEQI, and a low correlation was observed between DO and the six influencing factors, which may be attributable to the degradation characteristics of pollutants. Although some refractory pollutants are discharged into rivers by surface-runoff or point source pollution, resulting in high COD in the water, due to its refractory characteristics under natural conditions, the DO in water is not related to the total amount of pollutants.

**Table 8.** Analysis results of the relationships between the main elements exceeding standard limits and their influencing factors

| Indicators | DO | $\text{COD}_{\text{Mn}}$ | COD | $\text{BOD}_5$ | $\text{NH}_3\text{-N}$ | TP |
| --- | --- | --- | --- | --- | --- | --- |
| Application rate of chemical fertilizer per unit | 0.05 | 0.51 | 0.45 | 0.55 | 0.19 | 0.36 |
| Proportion of cultivated land irrigated area | 0.10 | 0.59 | 0.48 | 0.54 | 0.13 | 0.34 |
| Qw/Qs | 0.31 | 0.51 | 0.51 | 0.10 | 0.47 | 0.40 |
| Unit domestic waste removal and transportation | -0.03 | 0.45 | 0.33 | 0.18 | 0.51 | 0.37 |
| Ecological environmental quality index(EEQI) | -0.35 | -0.03 | -0.29 | 0.09 | 0.09 | 0.06 |
| Human activities quantitative index(HAQI) | -0.19 | 0.55 | 0.40 | 0.51 | 0.33 | 0.35 |

### Provincial Water Quality Improvement

Provincial boundary water quality is an important component of China’s surface water quality management, involving two or more provinces. The problems caused by provincial boundary water quality result in related contradictions between provinces. Therefore, the basic premise to improve the provincial boundary water quality is to analyze the water quality status of China’s provincial boundary water bodies and identify the key influencing factors. Upon analyzing the water quality of the 27 provinces, spatial differences in the water quality of different provinces in China were observed, and the evaluation results showed that the poor water quality was attributable to the DO, COD_Mn_, COD, BOD_5_, NH_3_-N, and TP exceeding the standard limits. In this study, 27 factors that possibly affect provincial boundary water quality were identified and integrated into six main factors by classification and index calculation, and the relationship between each factor and the water quality at provincial boundaries was analyzed. Based on the analysis results, it is necessary to focus on improving provincial boundary water quality in China by: 1. increasing awareness of environmental protection; 2. developing a clean economy; 3. controlling non-point source pollution; and 4. building an energy-saving and environment-friendly society.

## Conclusions

In this study, the water quality monitoring data from 540 provincial boundary monitoring stations of 27 provinces were analyzed in 2019, and the existing challenges in the water quality of China’s provincial boundaries were identified. The influencing factors that possibly affect the surface water quality were systematically and comprehensively analyzed, and the main factors leading to the deterioration of water quality at provincial boundaries were identified. The research results provide a strong scientific basis for the governance of water bodies at provincial boundaries in China. The main conclusions drawn are as follows:

1. The CCME-WQI shows that the composite index score ranges from 59.1– 97.2, a decrease from the excellent to marginal level. The single factor evaluation showed that provincial boundary water quality was dominated by Class **I** –**III** (accounting for 61.7%), and the water quality of inferior Class V accounted for 12.2%. The main elements exceeding standard limits were DO, COD_Mn_, COD, BOD_5_, NH_3_-N, and TP.
2. With respect to direct influencing factors, a negative correlation was observed between the water quality and application rate of chemical fertilizer per unit area (R^2^=0.15) and proportion of cultivated land irrigated area (R^2^=0.16) at provincial boundaries. There was no correlation between the water quality and total wastewater discharge/total surface water resources and unit domestic waste removal and transportation (R^2^<0.05).
3. There was no correlation between the CCME-WQI and the EEQI (R^2^ = 0.0112), and we analyzed the relationship between the six types of land use and CCME-WQI. The results showed that the R^2^ for farmland was 0.16 and 0.22 for building land.
4. We analyzed the relationship between the HAQI and the CCME-WQI: R^2^ was 0.1144, which had a more significant impact than EEQI. Considering the impact of social, economic, and cultural aspects on the aquatic environment, the correlation coefficients between these aspects and water quality were 0.1044, 0.1877, and 0.05, respectively.
5. COD_Mn_ and COD exhibited the lowest correlation with EEQI relative to the other five factors. BOD_5_ exhibited a strong correlation with the application rate of chemical fertilizer per unit, proportion of cultivated land irrigated area, and HAQI. NH_3_-N exhibited a strong correlation with Qw/Qs, unit domestic waste removal and transportation, and HAQI. There was no correlation between TP and EEQI.

## Acknowledgements

Jun Zhang provided overall guidance; Yuanyuan Gao and Shilu Zhang analysed the data; Mingxia Xu and Junyu He contributed analysis tools; Maoqing Duan wrote the paper.

## References

1. Qiao J., Liu F. (2014) Construction of System Framework of the Most Stringent Water Resources Management Regime. In: Qi E., Shen J., Dou R. (eds) Proceedings of 2013 4th International Asia Conference on Industrial Engineering and Management Innovation (IEMI2013).

2. Duan, M., Du, X., Peng, W., Zhang, S., Yan, L, 2019. A revised method of surface water quality evaluation based on background values and its application to samples collected in Heilongjiang Province, China. Water. 11(5), 1057–1073.

3. Maoqing Duan, Xia Du, Wenqi Peng, Cuiling Jiang, Shijie Zhang and Yang Ding, 2020. Necessity of Acknowledging Background Pollutants for Implement Sustainable Water Quality Management System in China. Science of total environmental.722, 137922.

4. Oliveira J C, Becegato V R, Barcarolli I F, et al, 2017. Environmental Characteristics and Water Quality of a Drainage Basin Impacted by Human Activities[J]. Environmental Management and Sustainable Development, 6(2):373.

5. Duan M, Du X, Peng W, et al, 2020. Quantitative assessment of background pollutants using a modified method in data-poor regions[J]. Environmental Monitoring and Assessment, 192(3).

6. Huo Z, Feng S, Kang S, et al, 2007. The Response of Water-Land Environment to Human Activities in Arid Minqin Oasis, Northwest China[J]. Arid Land Research and Management.

7. Liu, Hua Z, 2011. Current Situation and Main Pollution Sources of Rural Water Environment in China[J]. Advanced Materials Research, 281:113–116.

8. Sun Y, Chen G, Xu Z, 2020. Research progress of water environment, treatment and utilization in coal mining areas of China[J]. Meitan Xuebao/Journal of the China Coal Society, 45(1):304–316.

9. Savkin V M, Dvurechenskaya S Y, 2014. Resources-related and water-environmental problems of the integrated use of the Novosibirsk Reservoir[J]. Water Resources, 41(4):446–453.

10. Chen Z, Wu G, Wu Y, et al, 2020. Water Eco-Nexus Cycle System (WaterEcoNet) as a key solution for water shortage and water environment problems in urban areas[J]. Water Cycle, 1:71–77.

11. Liu X, Yua X, Yu K, 2012. The Current Situation and Sustainable Development of Water Resources in China[J]. Procedia Engineering, 28(none):522–526.

12. Leong E, 2013. Water Situation In China - Crisis Or Business As Usual?[J].

13. Dou M, Wang Y, 2017. The construction of a water rights system in China that is suited to the strictest water resources management system[J]. Water science & technology, 17(1-2):238–245.

14. China’s water resources annual report in 2018[J]. Haihe Water Resources(http://www.mwr.gov.cn/zzsc/tjgb/szygb/2018/mobile/index.html), 2019(4):43-43.

15. China Environment Statistical Yearbook in 2018 (http://www.stats.gov.cn/tjsj/ndsj/2018/indexch.htm).

16. Resource and Environment Science and Data Center of Chinese Academy of Sciences(http://www.resdc.cn/).

17. Environmental Quality Standards for Surface Water(GB2002-3838).

18. CCME, 1999. Canadian water quality guidelines for the protection of aquatic life: CCME Water Quality Index 1.0, Technical Report. In: Canadian Environmental Quality Guidelines, Canadian Council of Ministers of the Environment, Winnipeg.

19. de Rosemond S.; Duro DC.; Dubé M, 2009. Comparative analysis of regional water quality in Canada using the Water Quality Index. Environ. Monit. Assess. 156(1-4):223–240.

20. Hurley T.; Sadiq R.; Mazumder A, 2012. Adaptation and evaluation of the Canadian Council of Ministers of the Environment Water Quality Index (CCME WQI) for use as an effective tool to characterize drinking source water quality, Water Res, 46(11):3544–3552.

21. Zhigang Xu, Dafang Zhuang, Lin yang, 2009. Construction and Application of Regional Quantitative Model of Human Activity Intensity, JOURNAL OF GEO-INFORMATION SCIENCE,11(4):452–459.

22. Tappin A D, Rodriguez A N, Comber S D W, et al, 2018. The role of alkalinity in setting water quality metrics: phosphorus standards in United Kingdom rivers[J]. Environmental Science: Processes & Impacts, 20(10).

23. Albert E A, 2007. Chemical composition of drainage water around Kota Kinabalu City Centre area; Ammonia, nitrate and nitrite[J].

24. Rhew, Doughee, Soonju, et al, 2016. Relationships between water quality parameters in rivers and lakes: BOD5, COD, NBOPs, and TOC[J]. Environmental Monitoring and Assessment: An International Journal, 188(4):252.1.

25. Wang J, Lin Y, Zhai T, et al, 2018. The role of human activity in decreasing ecologically sound land use in China[J]. Land Degradation & Development.

26. Liu Y Y, Li Z H, Liang X J, et al, 2013. Assessment on Water Environment Quality of Jilongshan Sea Area in Zhanjiang in Winter[J]. Advanced Materials Research, 610-613:884–887.

27. Yang Y, Bao J, Song B, et al, 2020. Study on the Improvement Path of Water Environment Quality -- A Case Study of Tianjin[J]. IOP Conference Series: Earth and Environmental Science, 450(1):012126 (4pp).

